# Modeling Small Non-canonical RNA Motifs with the Rosetta FARFAR Server

**DOI:** 10.1101/031278

**Authors:** Joseph D. Yesselman, Rhiju Das

**Author notes:** Correspondence to: Rhiju Das.

## Abstract

Non-canonical RNA motifs help define the vast complexity of RNA structure and function, and in many cases, these loops and junctions are on the order of only 10 nucleotides in size. Unfortunately, despite their small size, there is no reliable method to determine the ensemble of lowest energy structures of junctions and loops at atomic accuracy. This chapter outlines straightforward protocols using a webserver for Rosetta Fragment Assembly of RNA with Full Atom Refinement (FARFAR) (http://rosie.rosettacommons.org/rna_denovo/submit) to model the 3D structure of small non-canonical RNA motifs for use in visualizing motifs and for further refinement or filtering with experimental data such as NMR chemical shifts.

## 1. Introduction

RNA plays critical roles in all living systems through its ability to adopt complex 3D structures and perform chemical catalysis (1). RNA structure appears highly modular in nature, defined through base pairing interactions. Nucleotides can either form highly structured helices composed of canonical Watson-Crick base pairs or small unpaired or non-canonical base paired regions in the form of junctions and loops (motifs) (2–4). Helices are, for the most part, structurally similar to each other, leaving non-canonical motifs to define the vast complexity of RNA structure and function. These non-canonical elements define the topology of the 3D structure of RNA by orienting the helices to which they connect and by forming long-range tertiary contacts that can lock specific global RNA conformations in place. In addition to defining the overall 3D structure of RNA (5,6), non-canonical motifs are the sites of small molecule binding and chemical catalysis (7–10). Many non-canonical motifs are on the order of only 10 nucleotides in size. Unfortunately, despite their small size, there is no reliable method to determine the ensemble of lowest energy structures of junctions and loops at near atomic accuracy. Nevertheless, to model RNA at high resolution, it is critical to achieve accurate solutions for these small motifs.

When their structures are solved experimentally, most motifs turn out to form complex arrangements of non Watson-Crick hydrogen bonds and a wide range of backbone conformations. Due to the large number of interactions possible and each nucleotide’s many degrees of internal freedom, it remains difficult to determine the lowest energy conformation (11). Recently, Fragment Assembly of RNA with Full Atom Refinement (FARFAR) was developed to help address this problem, FARFAR adapted the well-developed Rosetta framework for proteins to predict and design RNA non-canonical motifs (12). Out of a 32-target test set, 14 cases gave at least one out of five models that were better than 2.0 Å all-heavy-atom RMSD to the experimentally observed structure. While not perfect, this level of accuracy can be combined with even sparse experimental data, such as ^1^H chemical shifts, to obtain high confidence structural models, as was demonstrated recently in blind predictions with the CS-ROSETTA-RNA method (13). The motif models can also form building blocks for modeling more complex RNAs and has been tested in the RNA Puzzles trials (14). Our FARFAR method for large RNAs with complete folds has been reviewed recently (15). The current bottleneck for some of these motifs and for larger RNAs is the difficulty of complete conformational sampling (11). On-going work with stepwise assembly (SWA) attempts to resolve this issue (16), but this more advanced procedure requires greater computational expense and a complex workflow that is not yet straightforward to implement on a public server, except in the special case of refinement one-nucleotide-at-a-time crystallographic refinement (17). Stepwise assembly is available in the main Rosetta codebase, but will not be further discussed here.

This chapter outlines straightforward protocols that should enable anyone to use FARFAR through a simple web server. FARFAR (RNA De Novo) is part of the Rosetta Online Server that Includes Everyone (ROSIE) software, a push to give wide access to the algorithms found in the Rosetta 3.x framework (18). The web server requires no initial setup for the user; all that is needed is to supply a sequence and an optional secondary structure to obtain all-atom models of an RNA motif of interest.

### 1.1 FARFAR Calculation

The FARFAR structure-modeling algorithm is based on two discrete steps. First, the RNA is assembled using 1 to 3 nucleotide fragments from existing RNA crystal structures whose sequences match subsequences of the target RNA. Fragment Assembly of RNA (FARNA) uses a Monte Carlo process guided by a low-resolution knowledge-based energy function (19). Afterwards, these models can be further refined in an all-atom potential to yield more realistic structures with cleaner hydrogen bonds and fewer clashes; the resulting energies are also better at discriminating native-like conformations from non-native conformations. The two-stage protocol is called Fragment Assembly of RNA with Full Atom Refinement (FARFAR).

## 2. Materials

FARFAR (RNA De Novo) is a webserver implementation of the Rosetta RNA fragment assembly algorithm server using the ROSIE framework. ROSIE is a web front-end for Rosetta 3 software suite, which provides experimentally tested and rapidly evolving tools for the high resolution 3D modeling of nucleic acids, proteins, and other biopolymers. FARFAR (RNA De Novo) can be reached using any of the standard web browsers such as Apple Safari, Microsoft Internet Explorer, Mozilla Firefox and Google Chrome here: http://rosie.rosettacommons.org/rna_denovo/submit. [**Note to the editor - - address will change]**

## 3. Methods

This protocol outlines the steps to use the FARFAR (RNA De Novo) webserver located on the ROSIE website. Although it is possible to submit jobs without creating an account, having an account yields numerous benefits, such as email alerts when jobs are finished, as well as the ability to create private jobs that are not visible to other users. It is highly recommended to create an account when first visiting ROSIE. In addition to the FARFAR webserver, ROSIE also hosts many other Rosetta based applications with a continuous stream of novel applications in development.

### 3.1.1 Main Page Form

This demonstration of FARFAR (RNA De Novo) uses the GCAA tetraloop; the whole structure was determined through NMR spectroscopy by *Jucker et al.* (PDB 1ZIH) (20). This tetraloop has a sequence of gggcgcaagccu and secondary structure of `((((....))))` in dot parentheses notation (**Figure 1**). **Figure 2** shows the main submission form for the RNA De Novo server. The only required input is the sequence, from 5′ to 3′. This is typically in lower-case letters, but upper-case are acceptable and will be converted. Use a space, *, or **+** between strands. Note that this sequence is treated as RNA so that any Ts that appear in the sequence are automatically converted to Us for the calculation. Next, enter the secondary structure, in dot-parentheses notation. This is optional for single-stranded motifs, but required for multi-strand motifs. Note that even if a location is ‘unpaired’ in the input secondary structure (given by a dot, ‘.’), it is not forced to remain unpaired. Although this is optional for single stranded motifs, the results improve with the addition of the correct secondary structure. If uncertain about the secondary structure, consider utilizing the Vienna RNAfold webserver (21) (http://rna.tbi.univie.ac.at/cgi-bin/RNAfold.cgi) or other utilities described in this book. Alternatively, use chemical mapping techniques to estimate the structure; these methods have been recently tested in blind trials for their accuracy (22,23). In addition, note that there is currently a size submission limit of 32 nucleotides for FARFAR (RNA De Novo), as the amount of computation greatly increases as a function of number of residues.

**Figure 1:**
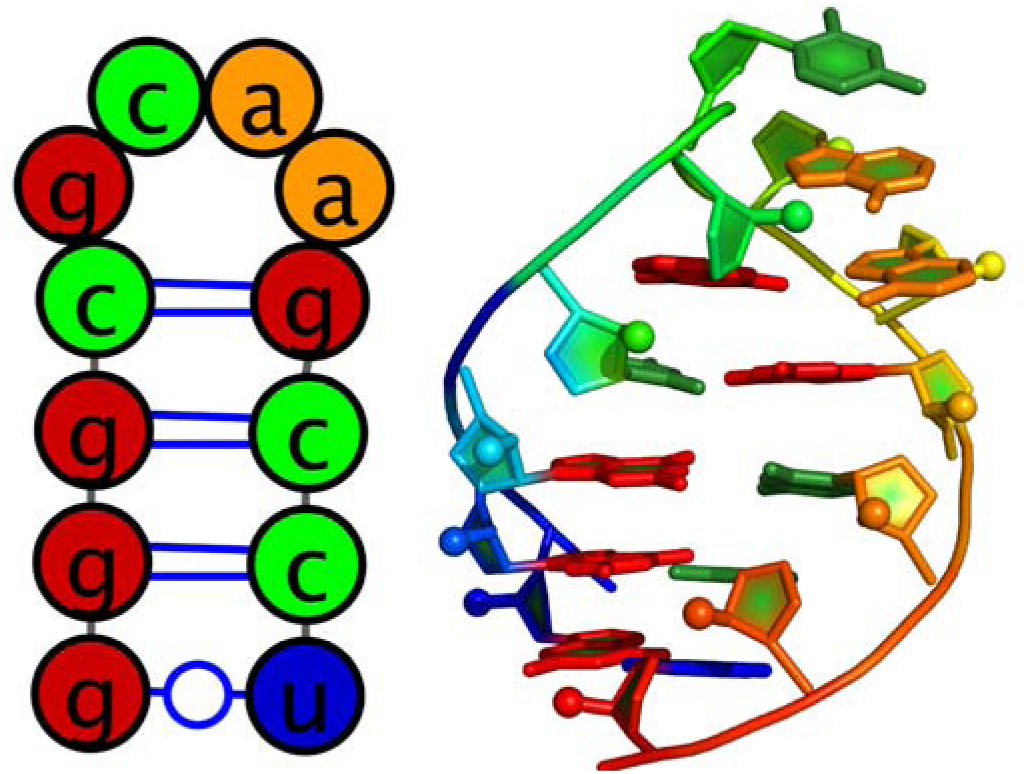
(Left) secondary structure of GCAA tetraloop. (Right) 3D structure of GCAA tetraloop (PDB: 1ZIH).

**Figure 2:**
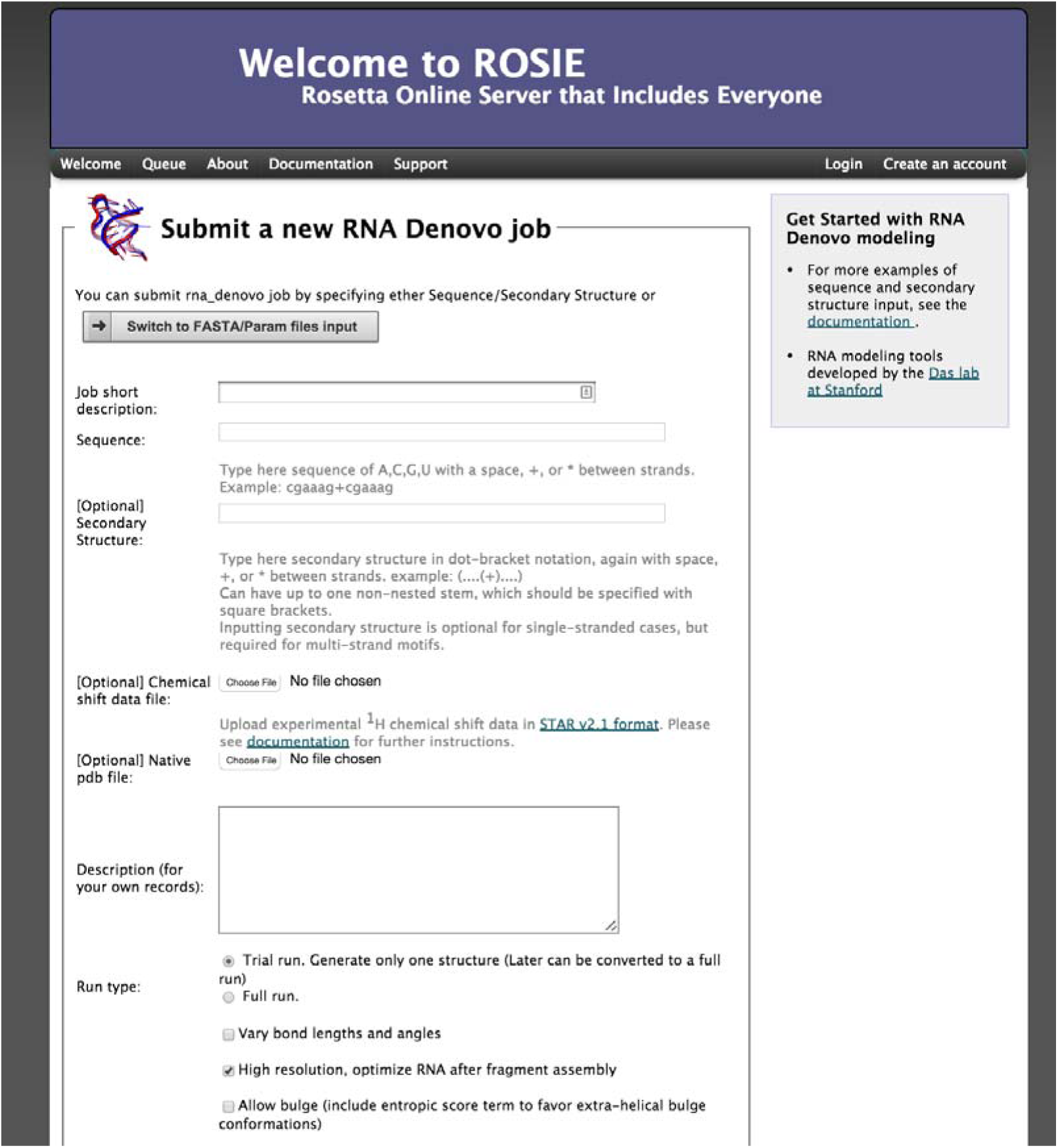
Main page of the FARFAR (RNA De Novo) webserver. Here the user can enter a sequence and secondary to submit a job to generation an all atom model of their construct.

There are two more optional arguments. First is a file containing the ^1^H chemical shifts determined by NMR spectroscopy. The format of this file follows the STAR v2.1 format used by the Biological Magnetic Resonance Data Bank (BMRB) (24). An example of the format is displayed in **Figure 3** with an explanation of each column. In addition, it is also possible to supply a native structure for the server to compare to for rmsd calculations. This file must be in PDB format, and for this case it is possible to download the structure from http://pdb.org/pdb/explore/explore.do?structureId=1zih. To supply a native structure, click the “Choose file” button next to native PDB-formatted file and select the appropriate file from your local hard drive.

**Figure 3:**
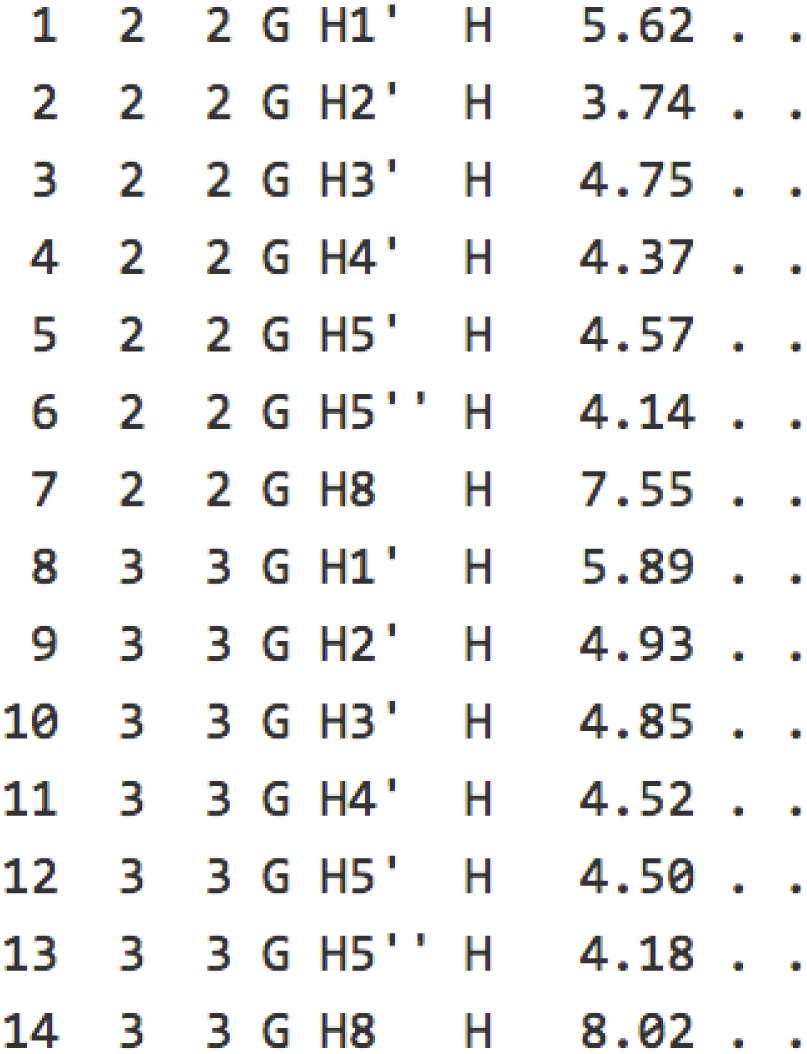
Example chemical shift data. Column description is as follows. 1) Atom entry number. 2) Residue author sequence code. 3) Residue sequence code. 4) Residue label. 5) Atom name. 6) Atom type. 7) Chemical shift value. 8) Chemical shift value error. 9) Chemical shift ambiguity code.

There are two ways of running a FARFAR (RNA De Novo) job. The first is a trial run, which generates only one structure with a limited number of fragment assembly steps. This is for testing purposes only, and allows you to confirm that the job is set up properly. The second is a full run that takes more computational time to complete and produces thousands of models. It is advised when setting up a job for a new sequence and secondary structure to always run the job as a trial. Then, using <u>PyMOL</u> or your favorite viewer, open the PDB file. This is particularly important if you have a multi-stranded motif - check that the strands are separated, and that any specified Watson-Crick pairs are reasonably paired. Once this is set up, go to the bottom of the page and click “Submit FARFAR (RNA De Novo) job”. Upon submission, a temporary status page will load (**Figure 4**).

**Figure 4:**
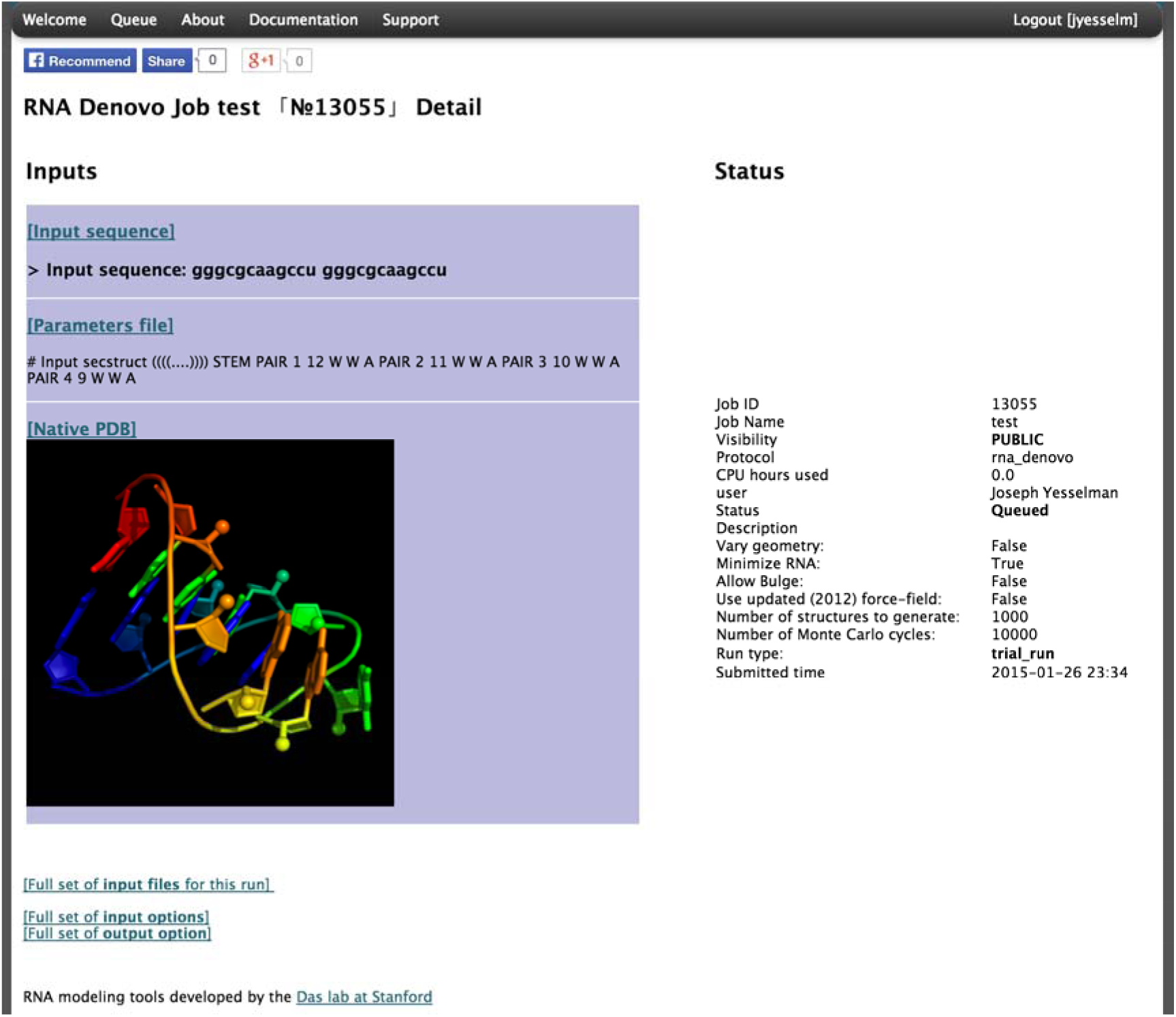
The status page for a submitted FARFAR (RNA De Novo) job.

### 3.1.2 Advanced Options

In addition to the options discussed above, there are a few additional options that may be used occasionally. First is “Vary bond lengths and angles”; typically each residue has a set of bond lengths and angles between atoms that are based on idealized parameters. Checking this option will allow these parameters to vary slightly based on the Rosetta force field energy. This can increase conformational search space if you are interested in a specific interaction between residues and was used in previous benchmark studies, but requires more computational time (12).

When checked, “High resolution, optimize RNA after fragment assembly” will perform the all-atom refinement after fragment assembly; it is not recommended to uncheck this unless you are interested in quickly seeing the initial results or would like to perform your own high resolution optimization. “Allow bulge (include entropic score term to favor extra-helical bulge conformations)”, will include conformations with residues bulged out and not interacting with other residues. If a residue is known to be extruded from the helix, this might be a good option to try to reduce the conformational space searched. When “Allow bulge (include entropic score term to favor extra-helical bulge conformations)” is checked, please note that residues that are bulged out will not be present in the final pdb model. “Number of structures to generate”, will change the number of final models, which will also greatly increase the time each run takes. “Number of Monte Carlo cycles”, controls the quality of each model; if models generated for a specific run have wildly different structures, then FARFAR has poor confidence in the accuracy (see next section). Increasing the number of Monte Carlo cycles can increase convergence, at the expense of greater computation.

### 3.3 Server Results

The server returns pictures of the best-scoring models from the five best-scoring clusters from the run in rank order by energy (**Figure 5**). The clustering radius is 2.0 Å by default. Click on the [Model-N] link to download the PDB file. The server returns cluster centers (without pictures) for the next 95 clusters as, as well as the top 20 lowest-energy structures. These may be valuable if you are filtering models based on experimental data. The server also returns a ‘scatter plot’ of the energies of all the models created. The x-axis is a distance measure from the native/reference model in RMSD (root mean-squared deviation) over all heavy atoms; if a reference model is not provided, then the RMSD is computed relative to the lowest energy model discovered by FARFAR. The y-axis is the score (energy) of the structure. In runs where a native structure is not supplied, the x-axis is a distance measure from the best scoring model found. As with nearly every Rosetta application, a hallmark of a successful run is convergence, visible as an energetic “funnel” of low-energy structures clustered around a single position. That is, near the lowest energy model there are additional models within ~ 2 Angstrom RMSD. In such runs, the lowest energy cluster centers have a reasonable chance of covering native-like structures for the motif, based on our benchmarks. A hallmark of an unsuccessful run is a lack of convergence - few structures within 2 Å RMSD of the lowest energy model. Below the scatter plot, there is a detailed table of all the score terms used to calculate the final score as well as the RMSD to the native structure (if supplied). A description of the meaning of each term can be found in **Table 1**.

**Figure 5:**
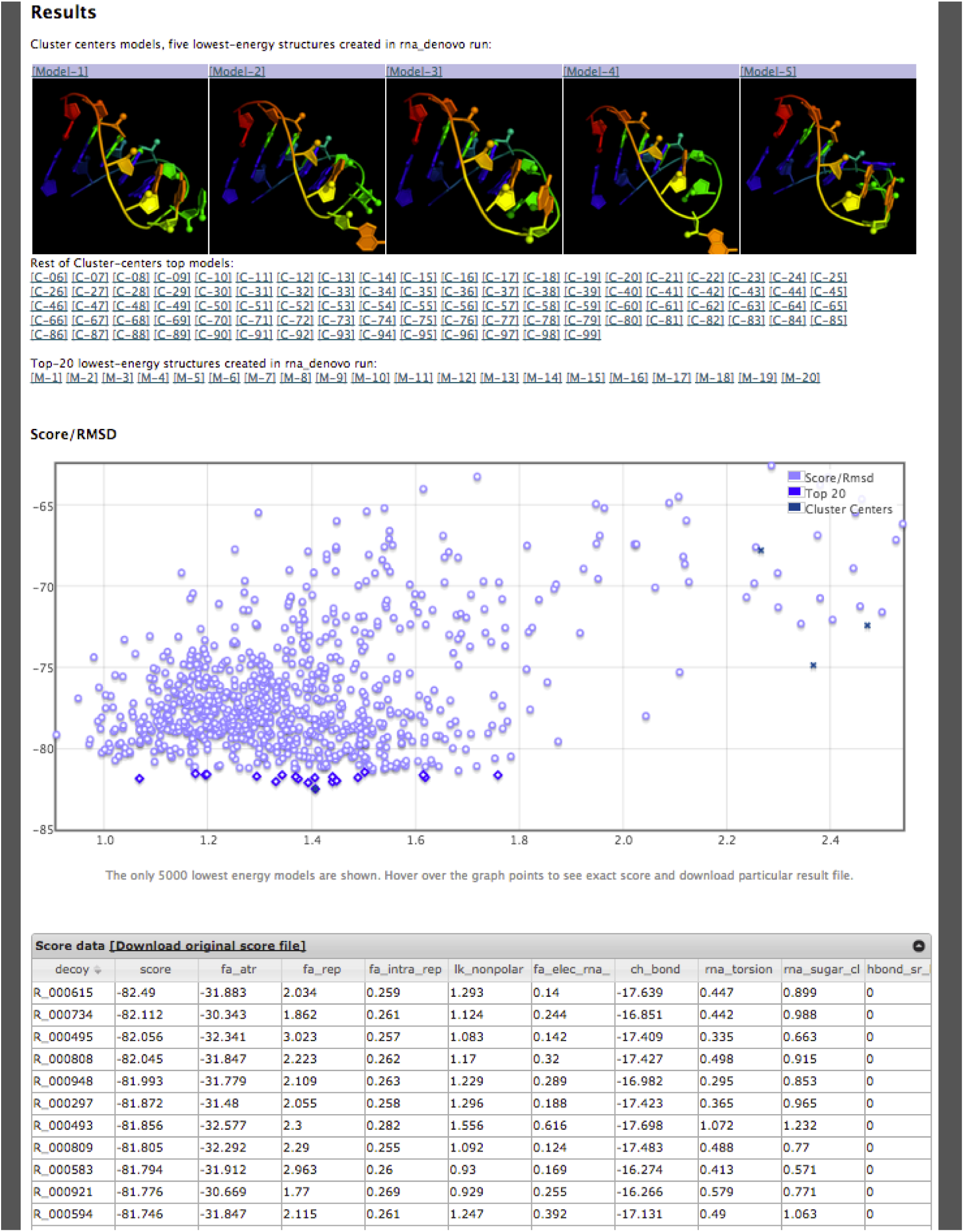
Results page for a RNA Denovo job.

**Table 1:** Score terms reported on RNA Denovo results page.

| Term | Definition |
| --- | --- |
| Score | Final total score |
| fa_atr | Lennard-jones attractive between atoms in different residues |
| fa_rep | Lennard-jones repulsive between atoms in different residues |
| fa_intra_rep | Lennard-jones repulsive between atoms in the same residue |
| lk_nonpolar | Lazaridis-karplus solvation energy, over nonpolar atoms |
| hack_elec_rna_phos_phos | Simple electrostatic repulsion term between phosphates |
| ch_bond | Carbon hydrogen bonds |
| rna_torsion | RNA torsional potential |
| rna_sugar_close | Term that ensures that ribose rings stay closed during refinement |
| hbond_sr_bb_sc | Backbone-sidechain hbonds close in primary sequence |
| hbond_lr_bb_sc | Backbone-sidechain hbonds distant in primary sequence |
| hbond_sc | Sidechain-sidechain hydrogen bond energy |
| linear_chainbreak | For 'temporary' chainbreaks, penalty term that keeps chainbreaks closed |
| N_WC | Number of Watson-Crick base pairs |
| N_NWC | Number of non-Watson-Crick base pairs |
| N_BS | Number of base stacks |
| Following are provided if the user gives a native structure |  |
| rms | All-heavy-atom RMSD to the native structure |
| rms_stem | All-heavy-atom RMSD to helical segments in the native structure |
| f_natWC | Fraction of native Watson-Crick base pairs recovered |
| f_natNWC | Fraction of native non-Watson-Crick base pairs recovered |
| f_natBP | Fraction of native base pairs recovered |

Visual representation of convergence of the models generated by FARFAR (RNA De Novo) can be found in **Figure 6**. As the figure demonstrates, there is high convergence in the top models found throughout the run. In addition, if one has ^1^H chemical shift data as mentioned it can also be supplied; this can greatly increase the convergence and accuracy of an FARFAR prediction run. This can be demonstrated through a simple GA tandem motif, first generating models without ^1^H chemical shift data (**Figure 6D**) yields the correct overall fold of the structure while incorrectly predicting the GA base pairs to be sheared instead of forming hydrogen bonds through their Watson and Crick edge (25). The ^1^H chemical shift data adds sufficient restraints to resolve the base pairing discrepancy, with all top 20 models having the correct base pairing as the NMR solved structure. Both the native pdb and the chemical shift file can be downloaded from http://rosie.rosettacommons.org/documentation/rna_denovo.

**Figure 6:**
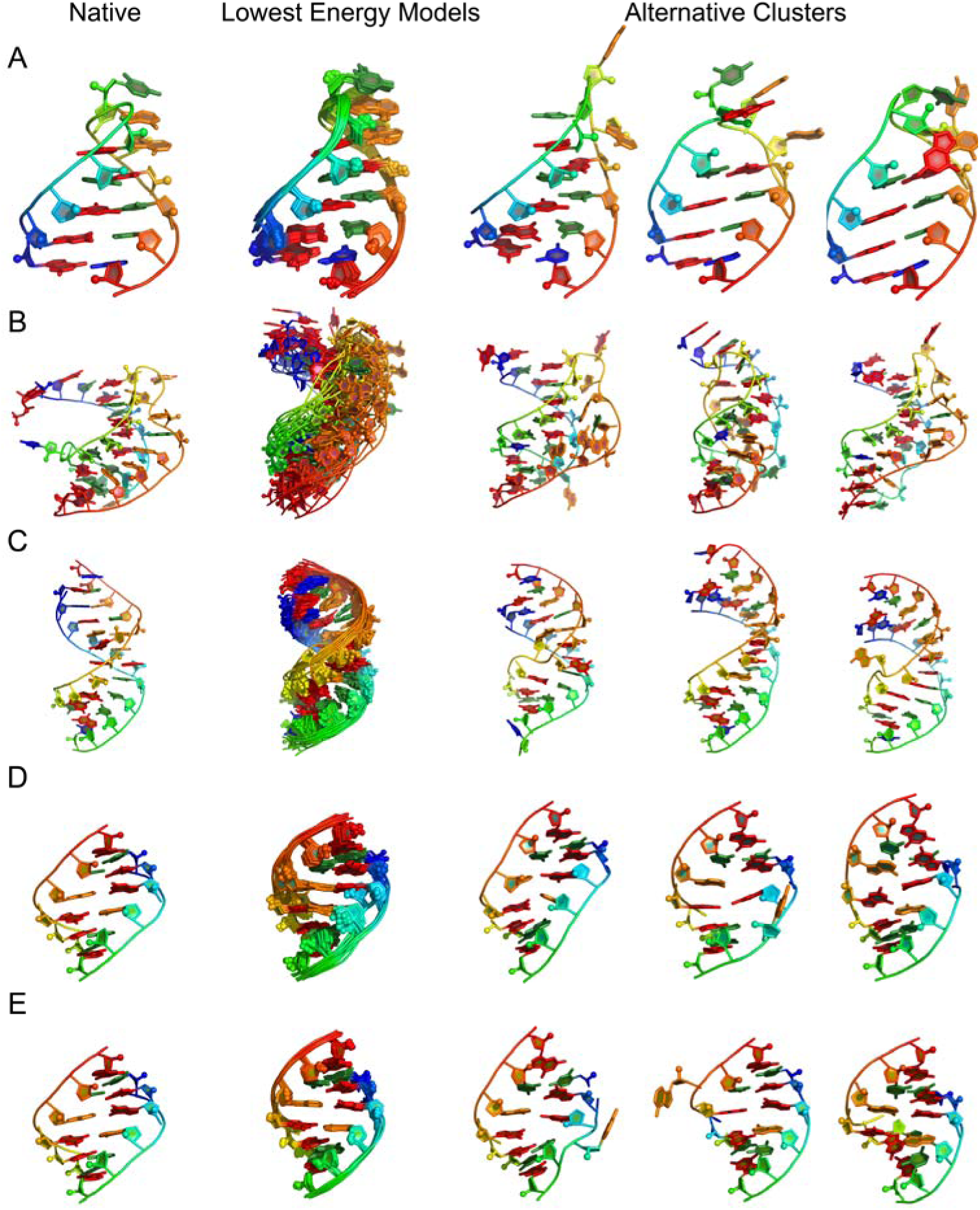
A) GCAA tetraloop (1ZIH), RNA Denovo lowest energy models displays a high level of convergence. B) Pseudoknot (1L2X) (26), less converged then tetraloop but also larger, still within 3Å heavy-atom rmsd for top model. C) 4×4 internal loop solved by NMR at PDB ID 2L8F (27), converges despite presenting 4 non-canonical base pairs. D) Tandem GA (1MIS) (25) without application of ^1^H chemical shifts. E) Tandem GA with ^1^H chemical shifts, demonstrates the improved convergence with the addition of ^1^H chemical shifts.

## Acknowledgments

We thank Sergey Lyskov for his thoughtful discussions and expert assistance with the ROSIE platform development. Writing of this work was supported by a Burroughs-Wellcome Foundation Career Award and National Institutes of Health Grant R01 R01GM100953.

